# Single-chain nanobody inhibition of Notch and avidity enhancement utilizing the β-pore forming toxin Aerolysin

**DOI:** 10.1101/2024.10.22.617501

**Authors:** Andrew C.D. Lemmex, Jeremy Allred, Jason Ostergard, Jake Trask, Hannah N. Bui, Michael J. M. Anderson, Benjamin Kopp, Oakley Streeter, Adam T. Smiley, Natalia S. Babilonia-Díaz, Bruce R. Blazar, LeAnn Higgins, Peter M. Gordon, Joseph M. Muretta, Wendy R. Gordon

**Author notes:** co-corresponding Authors. **Address Correspondence to:** Wendy R. Gordon and Joseph M. Muretta. equal contributions (Andrew C.D. Lemmex, Jeremy Allred).

## Abstract

Notch plays critical roles in developmental processes and disease pathogenesis, which has led to numerous efforts to modulate its function with small molecules and antibodies. Here we present a nanobody inhibitor of Notch signaling, derived from a synthetic phage-display library targeting the notch Negative Regulatory Region (NRR). The nanobody inhibits Notch signaling in a luciferase reporter assay and in Notch-dependent hematopoietic progenitor cell differentiation assay, despite a modest 19uM affinity for Notch. We addressed the low affinity by fusion to a membrane-associating domain derived from the β-Pore forming toxin Aerolysin, resulting in a significantly improved IC50 for Notch inhibition. The nanobody-aerolysin fusion inhibits proliferation of T-ALL cell lines with similar efficacy to other Notch pathway inhibitors. Overall, this study reports the development of a Notch inhibitory antibody, and demonstrates a proof-of-concept for a generalizable strategy to increase the efficacy and potency of low-affinity antibody binders.

## Introduction

The Notch signaling pathway is a highly conserved and central cell signaling pathway in metazoans involved in the regulation of cell differentiation, control of tissue patterning, and maintenance of tissue homeostasis^1,2^. One such example of its role in cell differentiation is the critical part that Notch signaling plays the immune system, specifically regulating the development of T-cells from progenitor cells in the thymus ^3^. Given its role in controlling cell fate, dysregulation of Notch signaling drivers multiple diseases including CADASIL, Alagille syndrome, and various cancers such as T-cell acute lymphoblastic leukemia (T-ALL)^4^.

The molecular mechanisms underlying the Notch signaling pathway are well understood. Notch ligands expressed on the surface of cells interact with Notch receptors on adjacent cells. This ligand-receptor interaction results in a mechanical force being applied across the Notch receptor^5^, which exposes a proteolytic site in a subdomain of the Notch receptor called the negative regulatory region (NRR)^6,7^, resulting in cleavage of the ectodomain by metalloproteases belonging to the ADAM family. Following cleavage of the ectodomain, gamma-secretase cleaves the Notch intracellular domain (NICD) which then translocates to the nucleus and interacts and forms a transcriptional complex with CSL and Mam to drive downstream Notch target gene expression^8^. Given the importance of the Notch signaling pathway and the multitude of roles it plays, there is significant interest in developing modulators that can be used to manipulate Notch signaling in both therapeutic and research contexts.

Several commercially available small molecules non-specifically inhibit the activation of Notch signaling, primarily through inhibition of the proteases involved in cleavage of the Notch receptor. Some examples include broad spectrum metalloprotease inhibitors such as Batimistat ^9^ and gamma secretase inhibitors^10^. Multiple blocking antibodies have also been developed that are capable of inhibiting Notch signaling by blocking ligand binding or targeting the NRR. Of the NRR binding inhibitory antibodies, several have therapeutically relevant effects in treating Notch related cancers^11,12^.

While antibodies have become an area of significant interest for the development of therapeutic molecules, they suffer from some notable limitations and drawbacks, the most significant of which is their large size (<100kDa) which limits their tissue penetration and uptake in tumors ^13^. One emerging tool in the antigen binding space is that of nanobodies. Nanobodies are single heavy chain antibody fragments derived from the single chain camelid VHH antibody which have emerged as powerful protein engineering platforms due to their relatively small size (15kDa), ease of expression in eukaryotic and prokaryotic cell systems, and their high solubility, and stability. Nanobodies exhibit superior tumor penetration properties compared to traditional antibodies, and though not completely immunologically naive, exhibit reduced immunogenicity compared to other non-human antibodies ^14–16^.

The majority of anti-Notch inhibitors are IgGs and IgG fragments. We used phage-display to identify nanobodies capable of modulating Notch signaling identifying a clone (S7) that binds to the Notch1 NRR with low affinity, but is capable of inhibiting Notch signaling and altering differentiation of a Notch dependent iPSC hematopoietic progenitor model. To increase the effective potency of S7 without altering its mode of binding, we localized it to the cell surface by fusing it to the GPI anchor binding toxin Aerolysin. This significantly enhanced S7 dependent Notch inhibition. The S7-Aerolysin construct was capable of inhibiting the cell-cycle progression of a T-Cell Acute Lymphoblastic Leukemia (TALL) cell line. Taken together this work demonstrates the feasibility of nanobodies in modulating Notch signaling, and also demonstrates the utility of Aerolysin in enhancing the effect of what would normally be a relatively weakly binding nanobody, which could be leveraged to enhance many other antigen binders of interest.

## Materials and Methods

### Plasmids

Plasmids used in this work were generated using Gibson assembly. The pADL22c phagemid was used to construct the synthetic nanobody phage-display library. The S7 nanobody was expressed for purification in pET15. The GFP binder nanobody was expressed in pET28 and was a generous gift from Dr. Margaret Titus. The Notch1 NRR was expressed from a plasmid derived from pTT5 and contained an N-terminal 8xHis tag and a C-terminal Avi-tag.

### Cell lines

Cell lines used in this study include DH5α cells for plasmid preparation, the TG1 and XLGold E. coli strains used for phage-display nanobody library production and phage-display biopanning. NiCo21 (New England Biolabs) were used for expressing S7 and anti-GFP nanobodies. The 6E suspension HEK cell line (Canadian Research Council) was used for Notch1 expression.

### Protein expression

#### Nanobodies

A 5mL LB culture with ampicillin was inoculated from a glycerol stab of BL21 E. coli transformed with pET3a expression plasmid coding for the S7 nanobody. This 5mL culture was grown overnight at 37C in shaker incubator. Next day, 5mL of the overnight culture was used to inoculate 200mL LB with ampicillin, which was then incubated at 37C overnight with shaking. Next day, 3 liters of LB with ampicillin was prepared (1L per flask) and 50mL of the 200mL overnight culture was used to inoculate each liter. 1L cultures were incubated at 37C with shaking until reaching an O.D. between 1.2 and 1.4. 1L cultures were then transferred to a 4C cold room to cool for 1 hour. After cooling for 1 hour the 1 liter cultures were moved back to a shaker incubator, and IPTG was added to each culture at a final concentration of 1mM to induce expression. Cultures were then incubated with shaking overnight at 18C. Next day cultures were harvested by centrifugation at 4000xG for 30 minutes at 4C, and pellets were stored at -80 for later purification.

#### Notch1 NRR proteins

Hek293F were grown and transfected according to manufacturer protocol [expifectamine 293 protocol]. In short, cells were grown to a density of 3.0*10^6 cells/mL, transfected with 1ug DNA per mL culture volume and incubated overnight at 37C and 5%CO2 with shaking. Next day, enhancer reagents 1 and 2 were added to cultures, after which cultures were incubated with shaking at 37C, 5% CO2 for 5 days. Following 5 days, cultures were harvested by centrifugation at 4000xG for 30 minutes. Supernatant was collected and stored at -20 for purification. Cell Pellets were disposed of.

### Protein Purification

#### Nanobodies

Previously harvested pellets were resuspended in Lysis buffer (50mM Tris, 500mM NaCl, 30mM Imidazole pH – 8.0) and equal volumes were transferred to 50mL Falcon tubes. Resuspended pellets were centrifuged at 6000xG for 5 minutes. Supernatant was poured off and then pellets were again resuspended in lysis buffer to a final volume of 40mL. 1 tablet of pierce EDTA-free protease inhibitor was added to each resuspended pellet as well as a tip-full [Need to measure out actual mass of this] of DNAse I. Resuspended pellets were then sonicated 4 times, 1 minute each, with rests on ice between each sonication. Sonicated pellets were then clarified via centrifugation at 24,000xG for 45 minutes. During centrifugation a 5mL HisTrap HP Sepharose Nickel column was prepared on an AKTA start chromatography system by washing 5CV H2O, washing 3CV Lysis buffer, washing 5CV elution buffer (50mM Tris, 500mM NaCl, 500mM Imidazole pH – 8.0), then finally washing with 5CV Lysis buffer. Following centrifugation clarified lysate was collected and applied to the 5mL HisTrap column. Following binding column was washed with 20CV lysis buffer, then protein was eluted with a 20CV gradient elution. 4mL fractions were collected. Fraction samples were run on an SDS-PAGE gel, and fractions containing the greatest amount of eluted nanobody were pooled and dialyzed overnight in 50mM Tris, 500mM NaCl, 100mM Imidazole pH – 8.0. The next day an additional step-down dialysis for 4 hours to 50mM Tris, 500mM NaCl, 50mM Imidazole pH – 8.0 was carried out before an overnight dialysis into 50mM HEPES, 20mM NaCl, 2% triton pH 7.9 to prepare samples for Ion-exchange chromatography on a HiTrap SP HP strong cation exchange column. Next day samples were loaded onto an equilibrated HiTrap SP HP column, and protein was eluted with a gradient elution into 50mM HEPES, 1M NaCl, 2% triton x-100 pH 7.9. Fractions were run on SDS-PAGE gels to assess for presence of S7 nanobody. To remove the triton x-100, Fractions containing S7 were pooled, dialyzed into 50mM HEPES, 20mM NaCl pH 7.9, and loaded back onto an equilibrated HiTrap SP HP column. Bound protein was washed extensively with 50mM HEPES, 20mM NaCl pH 7.9 before being eluted with a single step elution in 50mM HEPES, 500mM NaCl pH 7.9. Samples were then concentrated and run on a Superdex 75 10/300GL size exclusion column into either 20mM Tris, 150mM NaCl pH 8.0 for Cell based assays, or 20mM HEPES, 150mM NaCl pH 7.5 for Crosslinking experiments. Fractions from the size exclusion run were collected and concentrated down before snap freezing in liquid nitrogen and stored at -80.

#### NRR

Conditioned media containing expressed NRR constructs was brought to neutral pH and had additional NaCl added to a concentration of 500mM, along with addition of 30mM Imidazole. Conditioned media was then purified utilizing a HisTrap HP column manually loaded, washed and eluted. Wash buffer was 50mM Tris, 300mM NaCl, 5mM CaCl2 and 40mM Imidazole, pH 8.0. Protein was eluted in 50mM Tris, 300mM NaCl, 5mM CaCl2, 500mM Imidazole. Elution was collected and dialyzed overnight in 50mM Tris, 300mM NaCl, 5mM CaCl2. Next day protein was concentrated and purified via size exclusion chromatography on a Superdex 200 10/300GL column. Elution fractions were run on SDS-PAGE gel to assess for presence of purified NRR. Fractions containing NRR were then pooled, concentrated and snap frozen in liquid nitrogen. Samples were then stored at -80C.

### Nanobody phage-display library generation

The anti-GFP nanobody was synthesized as a double stranded DNA gBlock (IDT) and inserted into the pADL22c phagemid (Antibody Design Labs) using Gibson assembly. We used this plasmid as a template to construct a library of sequence variants with degenerate codons spanning the antigen recognition CDR loops. The library was constructed by multi-fragment overlapping PCR of three fragments. Fragment 1 was generated using primer pairs F1 and R1 that spanned the 5 prime portion of the nanobody through CRD1 with R1 containing degenerate codons across CDR1. Fragment 2 was generated with primer pair F2/R2, overlapped with Fragment 1 by 18 bp and spanned the 3 prime sequence of the nanobody. F2 introduced degenerate codons into CDR2 but left CDR3 intact. Fragment 1 and Fragment 2 were assembled into the pADL22c plasmid by Gibson assembly to generate a template plasmid with degenerate codons in CDR1 and CDR2 but not in CDR3. We used this plasmid library as a template to generate Fragment 3, which contained the 5 prime end of the nanobody to a region 5 prime to CRD3, spanning the degenerate CDR1 and CDR2 regions, and Fragment 4, which overlapped by 18 bp with Fragment 3 using primer pair F4/R4. F4 contained CDR3, thus introducing degenerate codons. For Fragment 4, we used two forward primers, the first contained 14 degenerate codons across the CDR3 loop, the second contained 18 degenerate codons. Fragments 3 and 4 were assembled by Gibson assembly into pADL22c. We transformed the resulting phagemid library into the TG1 E. coli strain and amplified nanobody-display phage with co-infection by M13 helper phage following established workflows outlined in ^17^.

### Phage-display Biopanning

Phage-display biopanning followed workflows outlined in Kontermann et al. ^17^ with several modifications. NRR protein was subjected to SDS-PAGE electrophoresis and transferred to nitrocellulose membrane. The NRR containing band was stained with ponceau, and the NRR band dissected in ∼0.5 cm square and then blocked with Phosphate Buffered Saline (PBS) containing 2% BSA. Blocked membranes were incubated for 1 hour with the phage display library diluted in PBS with 2% BSA and 0.1% Tween. Non-bound phage were removed and the membranes were washed extensively for 10 minutes with PBS with 0.05% Tween. Washing was repeated 10 times, and then the bound phage eluted by addition of 1 ml of 100 mM triethylamine, intubated for 10 minutes and transferred to microfuge tubes containing 0.5 ml 1 M Tris pH 7.5. The eluted phage were then incubated with 10 ml of log-phase TG1 cells, amplified with co-infection by M13 helper phage, and then purified by precipitation with 2.5 M NaCl 20% PEG prior to use in successive biopanning rounds.

### Conditioned media anti-NRR ELISA assays

Individual clones from e. coli infected with phage output from biopanning round 3 were diluted and plated on agar plates. Individual colonies were picked and used to inoculate 96-deepwell plates. Cultures were grown to OD 0.6 – 0.8 and induced. Next day plates were spun down and conditioned media was collected for use in the ELISA using HEK cell expressed Notch1 NRR as bait or tested for total nanobody expression. Clones that exhibited resilient NRR binding and expression at a single relative conditioned media dilution were evaluated for relative affinity by diluting the media in fresh media. 96-well immunosorbent plates [Nunc maxiSorp from fisher] were coated with NRR. Conditioned media containing the S7 nanobody was serially diluted across the NRR coated plates. Plates were washed and then incubated with an anti-HA antibody conjugated to HRP at a 1:1000 dilution. Plates were incubated for 1 hour at room temp and then washed 6 times with PBS. 100uL of TMB/H2O2 solution was added to each well and allowed to develop for 5 – 10 minutes. Reaction was stopped by adding equal volume 1N HCl to each well, and absorbance was measured on a plate reader at 450nm.

### Mammalian cell-line culture

U2OS cells were culture in 10cm dishes at 37C and 5% CO2. Cells were cultured in mCoy’s 5A (modified) media with 10% FBS from Thermo and passaged once reaching 90% confluency. Cells were passaged by aspirating media, washing 1x with PBS then adding 1mL Trypsin and incubating at 37C for 5 minutes. 9mL cell media was then added to quench trypsin reaction. Cells were passaged with a 1:4 dilution of cells to fresh media. For passaging of suspension cells--cells were grown to a density between 3.0*10^6 and 5.0*10^6 cells/mL. Upon reaching high density cells were counted and then diluted into fresh media for final seeding density of 5.0*10^5 cells/mL.

### Notch1 Signaling assay

Day before assay, U2OS cells are grown to confluence before passaging 1:1 with fresh media. Next day, 96-well half area plates are coated with 20ug/mL Notch ligand DLL4, which was resuspended to a stock concentration of 500ug/mL in PBS. Coated plates were incubated at 37°C for 1 hour. During the 1 hour plate incubation, U2OS cells were reverse transfected according to manufacturer protocol using lipofectamine 3000. Cells were co-transfected with 2ng/well Notch-Gal4 plasmid as well as 50ng/well Gal4::Luciferase and 1ng/well TK::Renilla Luciferase. The DLL4 coated plate was then washed 2×100uL PBS and transfected cells were plated on top. 3 – 6 hours post transfection cells were treated with relevant treatments by aspirating cell media and replacing with 80uL of media with appropriate treatment. Cells were then incubated overnight. Next day the Promega dual luciferase assay kit and protocol was used to read out signaling. In short, media was aspirated off of cells and cells were washed with 100uL PBS. 30uL Passive Lysis Buffer diluted with water was added to each well and incubated at room temperature for 20 minutes on an orbital shaker. Following incubation, 5uL of cell lysate was transferred from each well to a 96-well half-area assay plate. Signaling was then read-out in a Biotek Synergy H1 microplate reader with dual injectors. The ratio between the luciferase expression and constitutive Renilla luciferase luminescent was then calculated and plotted using PRISM.

### Induced Pluripotent Stem Cell Differentiation

Induced pluripotent stem cells were derived from random donors and generated using the Sendai virus technique. iPSCs were cultured on Geltrex™ (Thermo Fisher Scientific, A1413302) coated 6-well plates in mTeSR™1 (STEMCELL, 85850), On Day -1 a monolayer differentiation culture was generated by enzymatically dissociating iPSC colonies using TrypLE Select (Gibco, 12563-029) and plated on Geltrex coated 6 well plates at a density of 10,000 iPSCs/cm^2^ in 2ml/well of mTeSR1 + 10uM Y-27632 dihydrochloride (Tocris, 1254). Cells were cultured at 37°C, 5% CO_2_, 5% O_2_ for 24h. On Day 0, the media was then exchanged for 2ml/well of mesoderm induction media (MIM), composed of STEMdiff APEL 2 Medium (STEMCELL, 05275) + 50ng/ml Recombinant Human FGF-basic (Peprotech, 100-18B), 50ng/ml Human BMP-4 (E.coli derived) (Peprotech, 120-05ET), 15ng/ml Human/Murine/Rat Activin A (E.coli derived) (Peprotech, 120-14E) + 2mM LiCl (Sigma-Aldrich, 203637), and 1uM Y-27632 dihydrochloride. At D2, the media was then exchanged for 3ml/well of hemogenic endothelium media (HEM), composed of STEMdiff APEL2 + 50ng/ml Recombinant Human FGF-basic, 50ng/ml Human VEGF 165 (Peprotech, 100-20), 6uM SB-431542 (hydrate) (Cayman Chemical Company, 13031), and 3uM CHIR99021 (Sigma-Aldrich, SML1046). At Day 4, the current media was collected and cells were washed with 1x DPBS w/o Ca^2+^ & Mg^2+^ and dissociated in 1x TrypLE Select. They were then counted and replated at 41,700 cells/cm^2^ onto 12-well plates coated with irradiated mouse OP9-DLL4 stromal cells (kindly provided to our lab by Juan Carlos Zúñiga-Pflücker) in 1ml/well of EHT media, composed of aMEM (Thermo Scientific, 12000022; reconstituted per manufacturer’s instructions) + 10% FBS + 50ng/ml Recombinant Human SCF/c-kit (R&D Systems, PRD255-50), 50ng/ml Human TPO (Peprotech, 300-18), 50ng/ml IL-6 (Peprotech, 200-06), 10ng/ml Human IL-3 (Peprotech, 200-03), and 10ng/ml Human Flt3-Ligand (Peprotech, 300-19). Then, S7 at 2.5uM, 5uM and 10uM as well as 10uM anti-GFP, and equal volume buffer (20mM Tris pH7.5 150mM NaCl) control conditions were added in triplicate. Volumes added per well for each condition were normalized using the aforementioned buffer. At Day 6, an additional 0.5ml of the same respective media was added to each well. At Day 8, both non-adherent cells and adherent cell fractions (via TrypLE digestion) were collected and counted using a Cellaca MX High Speed Cell Counter (Nexcelom Bioscience, MX-AOPI) before preparing samples for analysis via flow cytometry.

### CL-MS

Purified S7 (20uM) was mixed with Notch1 NRR (10uM) with the addition of DSSO (1.2mM final concentration - 200uL final volume). Samples were mixed by gentle inversion and flicking of the tube followed by spinning down the solution in a table-top microcentrifuge. Samples were then incubated at 4C for 1 hour. The DSSO cross-linking reaction was quenched by the addition of 1M Tris pH 8.0 added to a final concentration 20mM. Protein was then precipitated via chloroform/methanol precipitation as previously described (^18,19^. In short, 0.8mL methanol was directly to the crosslinked-protein solution. Samples were vortexed briefly, then centrifuged for 20 seconds at 9000xG. Following centrifugation, 0.2mL of chloroform was added to the sample. The sample was then vortexed and centrifuged at 9000xG for 20 seconds. Following centrifugation, 0.6mL deionized water was added to the methanol/chloroform solution. Sample was vortexed for 5 seconds and centrifuged for 1 minute at 9000xG. The aqueous upper layer was carefully removed and discarded, being careful not to disturb the protein flocks visible as white flakes in the interphase. 0.6mL of methanol was then added and the sample was then vortexed followed by centrifuged for 2 minutes at 16,000xG. Supernatant was removed and the pellet was air dried. Prepared Pellets were then stored at -80. Precipitated protein was then trypsin digested and run through LC-MS^3^ and analyzed with proteome-discoverer 3.0 utilizing the Xlink plugin

## Results

### Identification of a candidate NRR binder

We designed a synthetic nanobody library to utilize in a phage-display biopanning strategy against recombinantly expressed Notch1-NRR (Fig1A). Following three rounds of selection, single colonies were picked and used to inoculate cultures in 96-deep-well plates to generate nanobody conditioned media. An amber stop codon encoded in the phage constructs results in secretion of the nanobody when expressed in e.coli. Media from each clone was then collected and used to perform ELISA assays with Notch1-NRR coated plates in order to screen for potential nanobody binders of interest. From these ELISA screens we identified several potential binders and selected the clone (strain 7, referred to as S7) that exhibited the highest binding for further testing and characterization (Fig1B). In follow-up ELISA Assays, a titration of dilutions of S7-conditioned media demonstrated specificity towards S7 but not BSA and other bait proteins (Fig1C). Lastly, in order to estimate the Kd of S7, we utilized microscale electrothermophoresis (MST). S7 was recombinantly expressed and purified from e.coli. Purified protein was titrated against 50nM of N1-NRR. Results from the assay indicate that S7 is binding the NRR with a kD of roughly 19.36uM (Fig1D). Taken together these data indicate that S7 is capable of binding to the NRR, though relatively weakly.

**Figure 1.**
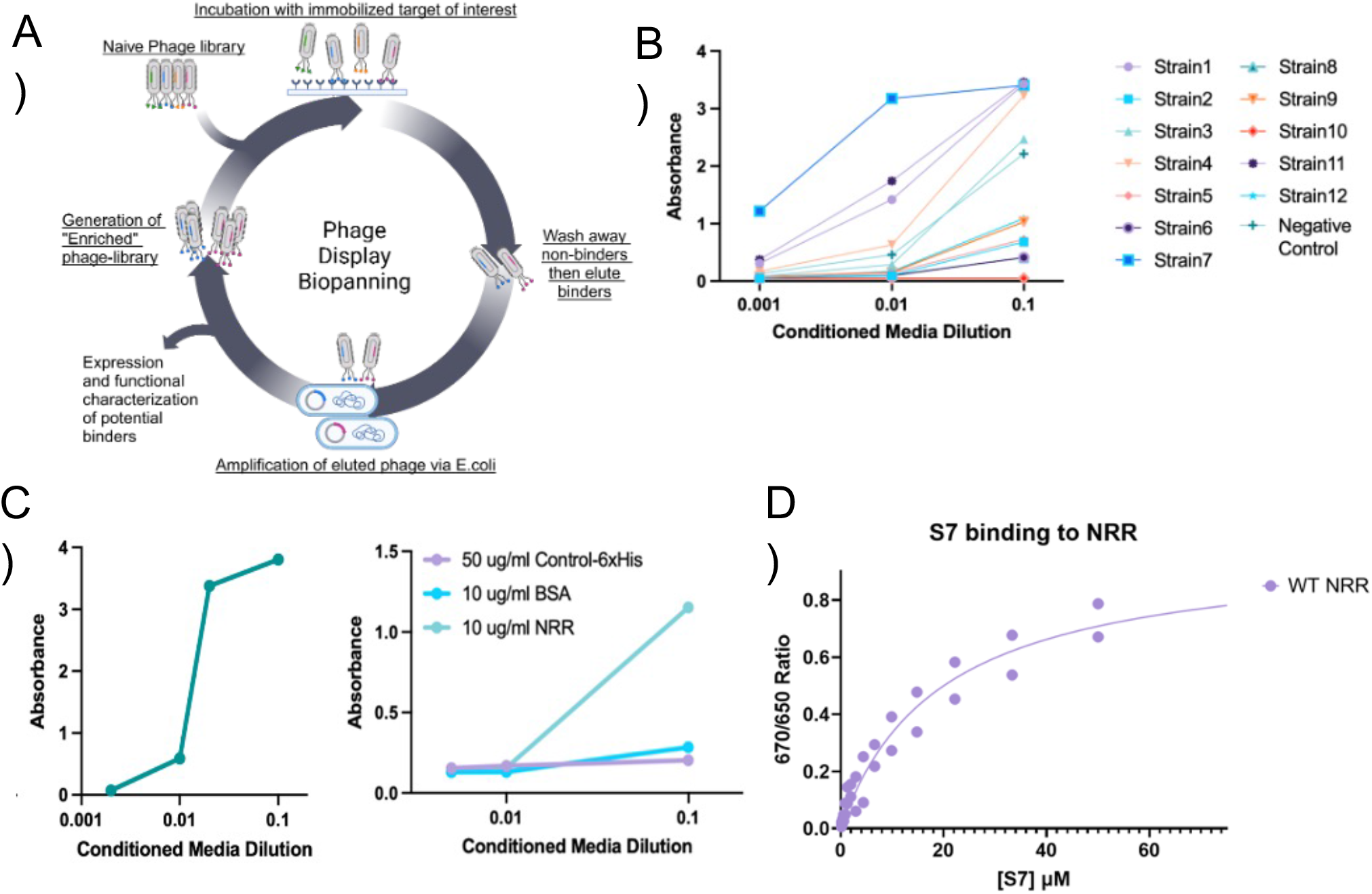
Identification of a Notch1 NRR binding Nanobody. (A) Anti-NRR ELISA of 12 candidates at 3 conditioned media dilutions. (B) Sanger sequencing of Strain 3, Strain 4, Strain 7, Strain 10, Strain 11, and Strain 12. Strain 10 contains an amber TAG codon which is read through in TG1 cells. (C) Testing of Strain 7 conditioned media on plates coated with an irrelevant protein M04-6xHis, BSA, or NRR. (D) Testing of Strain 7 conditioned media on plates coated with NRR or mClover3-NRR-mRuby3.

### S7 inhibits Notch activation in cell-based signaling assay

We next evaluated the impact of S7 on Notch1 signaling using an established Notch1 signaling assay. (Fig2A) provides a general overview of the process, but briefly, U2OS cells were co-transfected with a Notch1-gal4 expressing plasmid, a UAS-Firefly Luciferase plasmid and a pRL-TK plasmid expressing Renilla luciferase to allow for normalization of each condition for effects such as cell health, cell density, etc. Transfected cells were plated in triplicate in 96-well plates coated with or without the Notch1 ligand Dll4. Cells were then treated 3 hours after plating with either a Gamma Secretase inhibitor (GSI, Notch1 inhibition control), DMSO (vehicle control), cell media, or increasing concentrations of S7 nanobody. The next day, cells were prepared for reading out luciferase and renilla signals in a plate reader as described in the methods section. Cell lysate from each condition was reacted sequentially with Firefly and Renilla luciferases, and luminescence read out on a plate reader. Consistent with other studies, GSI treatment had comparable levels of Notch signaling to no ligand levels. The S7 titration demonstrated a dose dependent reduction in Notch1 signaling in the range of concentrations tested up to 40uM S7 (Fig2B). Follow-up dose responses of N1NRR to titrations of S7 demonstrated an apparent IC50 of 4.5uM (Fig2C). These data indicate that even though S7 is a weak binder of the NRR, it does inhibit activation of Notch signaling.

**Figure 2.**
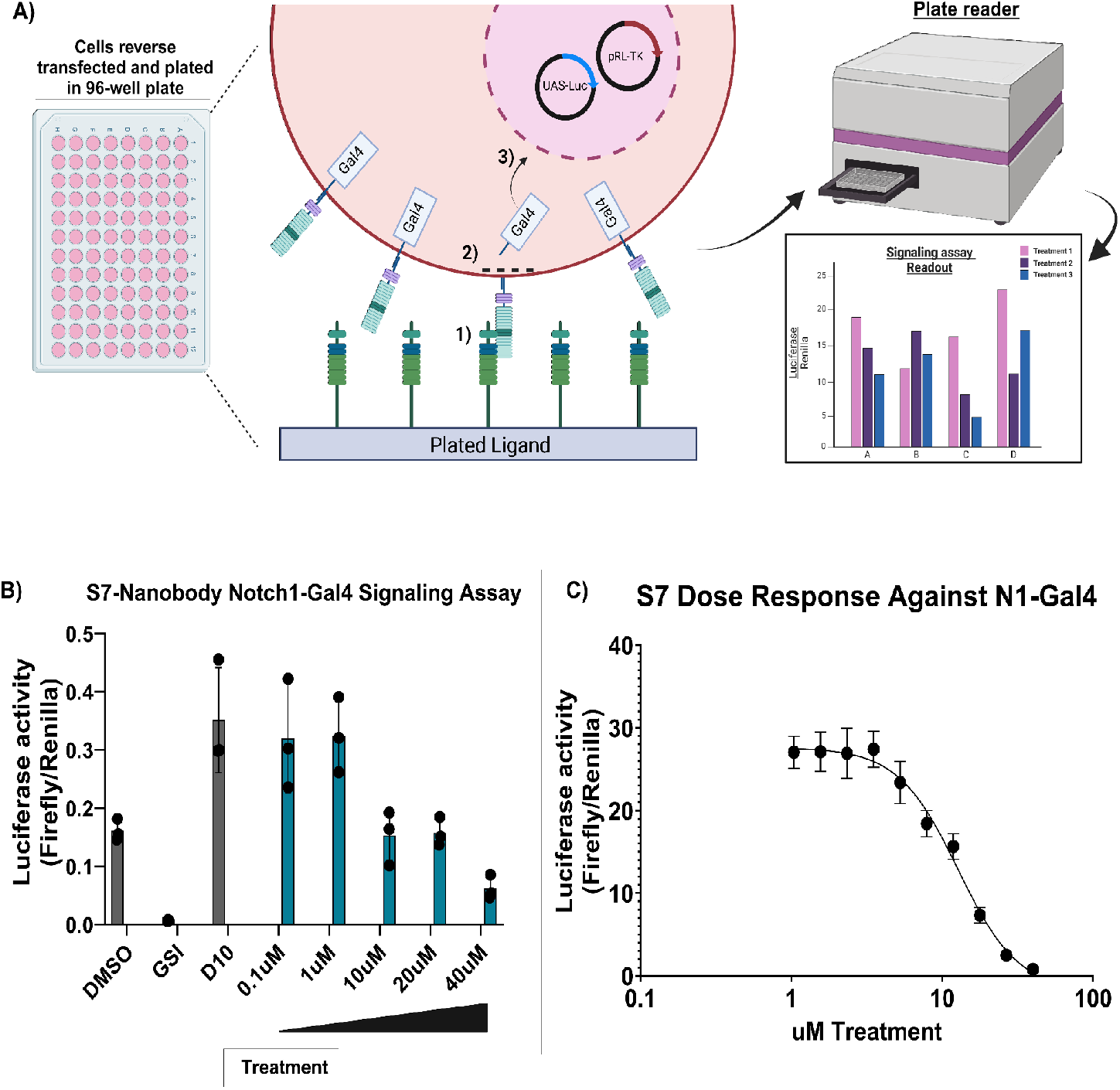
S7 inhibits Notch activation in cell-based assay. (A) Schematic of signaling assay. Cells transfected with N1-Gal4 promoter are plated in plates precoated with Notch ligand. The Receptor binds the Notch ligand (1), leading to S2 and S3 proteolytic cleavages, releasing the intracellular Gal4 domain (2) and translocation to the nucleus where it drives transcription of firefly luciferase (3). Firefly luciferase signal is normalized against constitutive expression of Renilla luciferase. (B) Effect of serial dilution of S7 on Notch1-Gal4 signaling. Each point represents 3 technical replicates

### S7 inhibits definitive hematopoiesis

While the cell signaling assay provided useful information in regards to the capability of S7 to modulate Notch signaling, it relies on the overexpression of the Notch receptor and may not accurately reflect the inhibitory capabilities of the S7 nanobody in more physiologically relevant contexts. To test the impact of S7 on endogenous Notch1 activity in a physiologically relevant context we evaluated S7’s effect on human hematopoietic progenitor cell differentiation from induced pluripotent stem cells (iPSCs). The differentiation of iPSCs into the more mature “definitive” hematopoietic lineages capable of generating multipotent progenitors is dependent by Notch1 activation.^20–22^ In the absence of Notch1 activation, iPSCs develop into lineage-restricted embryonic and primitive hematopoietic progenitors^23^. To drive Notch1-dependent definitive differentiation, iPSCs were plated on a monolayer of mouse OP9 cells engineered to display the Notch1 ligand DLL4 (OP9-DLL4) in addition to HSPC media for inducing differentiation into hematopoietic stem cells. Differentiation conditions are described in detail in Methods, with Figure 3A outlining the workflow for the assay. In short, iPSCs were first differentiated into hemogenic endothelial cells (Day 0-4). At that point, cells were collected and plated onto OP9-DL4 cells in hematopoietic specification media with varying amounts (10uM, 5uM or 2.5uM) of S7 nanobody, or corresponding volumes of S7 nanobody buffer without S7, or 10uM anti-GFP (the scaffold used to engineer S7) as negative controls. At Day 8 of differentiation, cells were collected and analyzed via flow cytometry for immunophenotypic markers of definitive (CD43+CD34+CD235/41a-) vs primitive (CD43+CD235a/41 differentiation. The overall production of hematopoietic progenitor cells (CD43+ HSPCs) was not statistically different between the buffer only and anti-GFP nanobody controls. S7 treatment resulted in a modest but significant increase in total CD43+ HSPCs across dose levels (Fig 3C) that likely represents increased CD43-CD73-cells successfully undergoing EHT at higher S7 concentrations or increased proliferation of primitive vesus definitive cells. Within the CD43+ subset, Notch1 inhibition via S7 resulted in a modest increase in the proportion of CD235/41a-primitive HSPCs (Fig 3B) but significantly increased absolute number of primitive HSPCs up to 5uM (Fig 3D). Conversely, Notch1 inhibition significantly decreased both the number (Fig 3E) and proportion (Fig 3F) of CD34+CD235/41-definitive HSPCs. The observed effect of S7 in reducing definitive HSPCs and increasing the ratio of final primitive HSPC production to definitive HSPCs is consistent with expectations of how a Notch1 inhibitor would impact differentiation of iPSCs to definitive HPSCs. These data also demonstrate that S7 is able to maintain inhibitory effects in physiologically relevant contexts despite its relatively weak binding.

**Figure 3.**
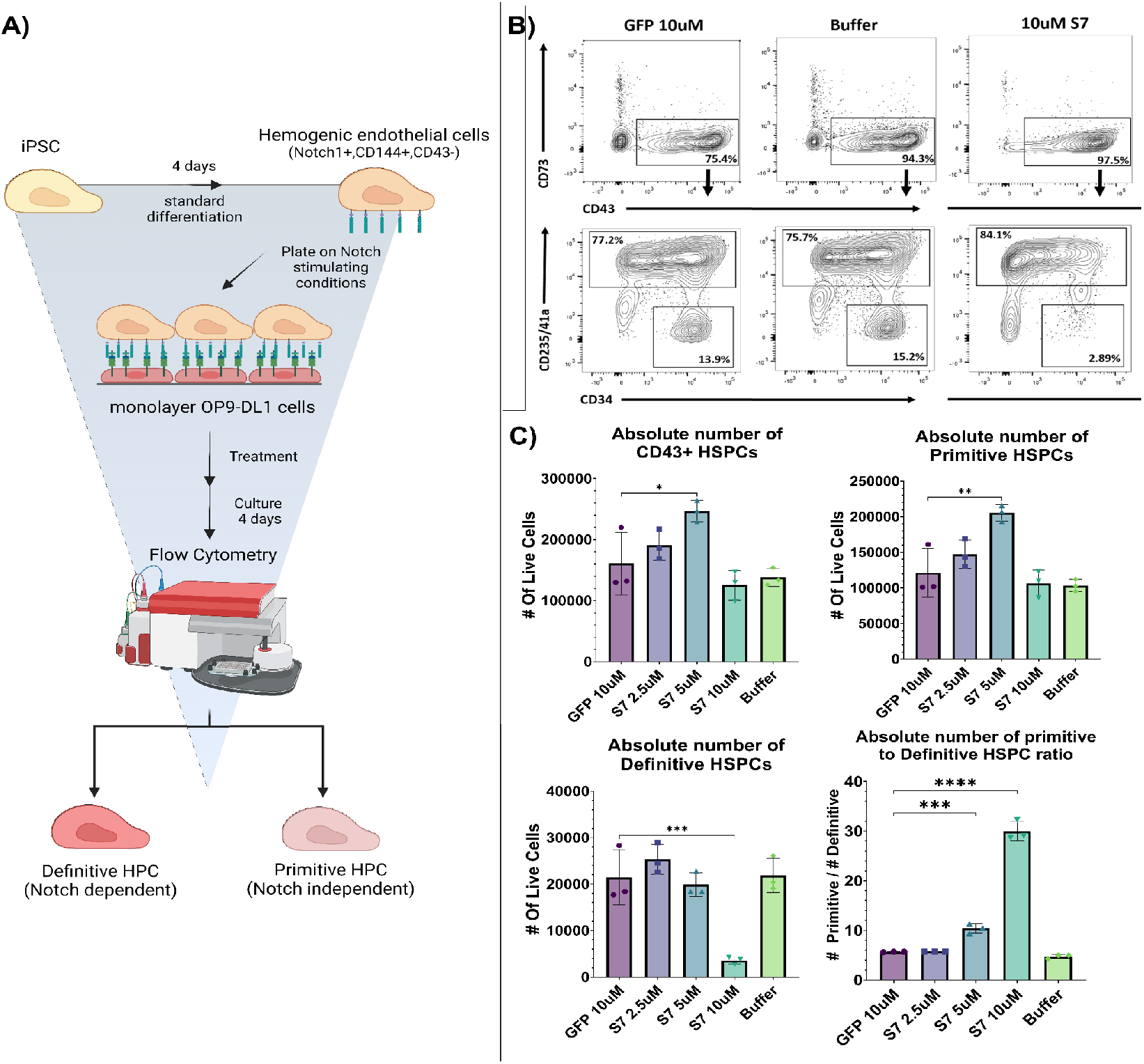
S7 inhibits differentiation of hemogenic endothelial cells into definitive hematopoietic cells. A. Assay scheme B. Flow cytometry contour plots showing relevant T-cell populations in the presence of buffer, GFP nanobody control, and S7 nanobody. C. Quantification of cell populations from panel B.

### S7 efficacy can be enhanced via fusion to the bacterial toxin Aerolysin

While S7 demonstrated inhibitory activity against Notch in both synthetic signaling assays as well as a physiologically relevant differentiation assay, its low affinity requires high concentrations of nanobody to observe any inhibitory effect. We reasoned that one way to address this issue would be to increase local concentrations of S7 by tethering it to a high-affinity cell-surface binder. To achieve this we chose the bacterial toxin Aerolysin as the fusion partner for S7 (Fig 4A). Aerolysin is a founding member of the pore forming toxin family known as Beta-Pore-Forming-Toxins (β-PFT), and is produced by the bacteria *Aeromonas hydrophila*. Aerolysin recognises and binds with high affinity to a highly conserved class of cell surface proteins known as Glycophosphatidylinositol-anchored proteins (GPI-AP). GPI-APs are cell-surface proteins that have been post-translationally modified via the addition of a Glycophosphatidylinositol moiety on the C-terminus of the protein^24,25^. Additionally, due to the significant structural studies undertaken to understand Aerolysin’s pore-forming process several mutants have been identified that arrest pore-formation but do not negatively impact binding^26^. One such mutant, Aerolysin T253/A300C, was selected as the fusion partner for S7. S7 was fused to the N-terminus of aerolysin and expressed and purified from e.coli as described in the methods. S7-aerolysin was initially evaluated in the Notch1-Gal4 signaling assay comparing a dose response of S7-aerolysin vs S7 nanobody. We found that S7-Aerolysin had significantly increased potency compared to S7, with a comparable inhibitory effect at 1uM with S7 at 20uM. The estimated IC50 of S7-aerolysin was 67.8 nM compared to an IC50 of 4.5uM for S7, a 67 times increase in potency (Fig 4B). These data show that Aerolysin provides a potent and simple method for increasing the efficacy of S7 without the need for further engineering of the nanobody.

**Figure 4.**
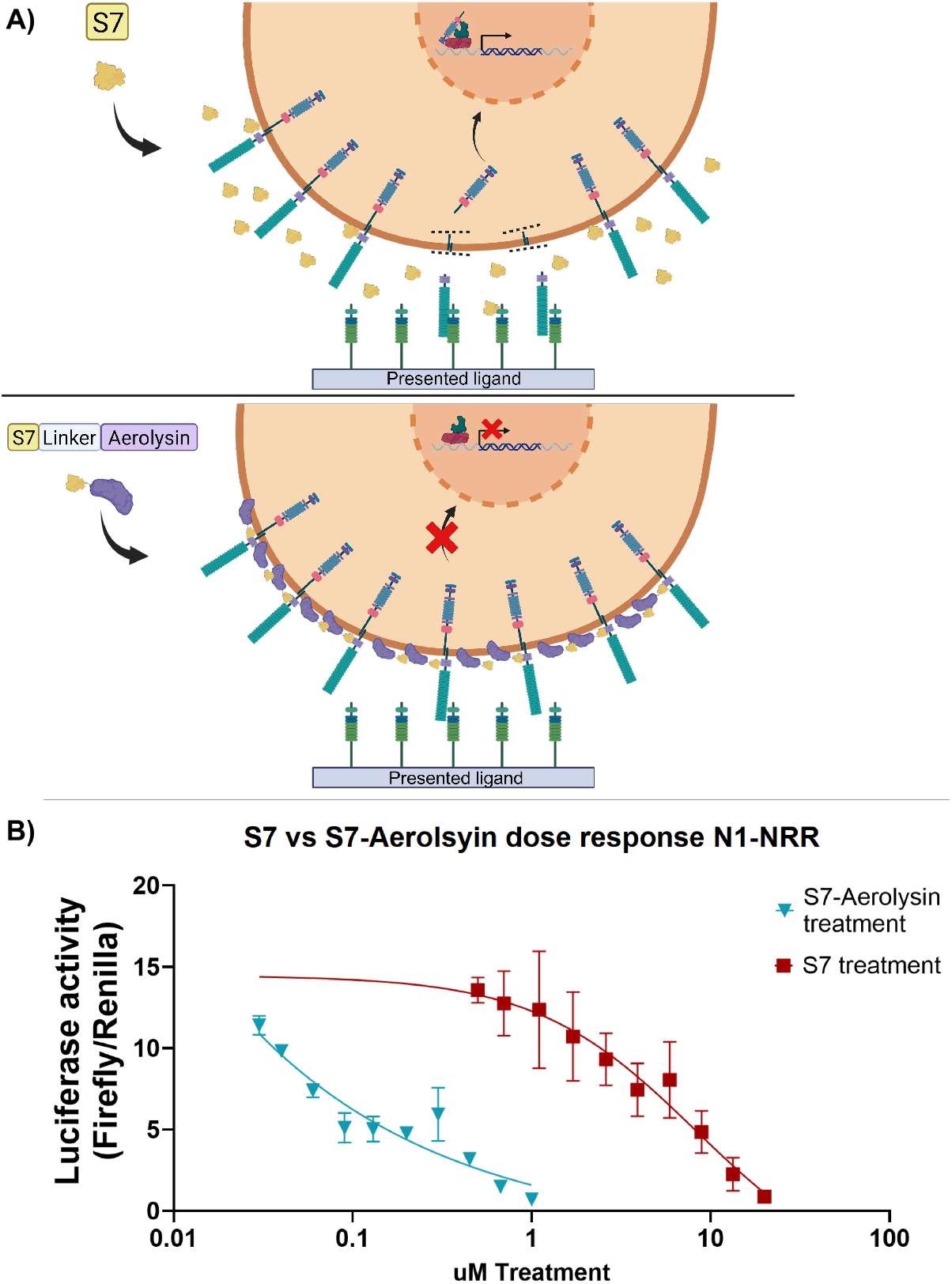
S7 fused to Aerolysin improves efficacy of S7 inhibitory effect. (A) Cartoon representation of S7 fused to the N-terminus of Aerolysin and the localization of S7 to the surface of the cell via Aerolysin. (B) Cartoon representation of the assay in C, in which plates are coated with a titration of DLL4 ligand, cells with the Notch reporter are plated and then treated with a fixed concentration of S7 or S7-aerolysin to determine if the treatments shift the response to DLL4. (C) The response of a Notch reporter of activation in a cell-based signaling assay to treatment with either S7 or S7-Aerolysin at different concentrations. (D) The response of U2OS cells in the signaling assay plated on different concentrations of DLL4 ligand while treat with either 10uM S7 or 250nM S7-Aerolysin.

### S7-Aerolysin uniquely affects surface Notch

While one explanation for the increased potency of the S7-Aerolysin fusion compared to S7 is that Aerolysin simply localizes more S7 on the surface of the cell, near the Notch receptor, creating a higher local concentration/avidity effect, an alternative explanation could be that S7-Aerolysin uniquely interacts with and depletes the Notch receptor on the surface of the protein. This could be due to activation of alternative cell membrane recycling pathways, which aerolysin is known to activate in certain contexts. To assess effects of S7-aerolysin on cell surface expression of Notch, we used flow cytometry to quantify cell-surface Notch using a doxycycline inducible U2OS cell line which expresses a FLAG-Notch1-GFP construct. Cells were plated in 24 or 48-well plates and expression of the FLAG-Notch1-GFP construct was induced with 1ug/mL doxycycline. 3 hours later, various treatments were added and expression of FLAG-Notch1-GFP was maintained with 1ug/mL doxycycline. Cells were allowed to incubate overnight. The next day, cells were washed with PBS and then incubated with an Anti-FLAG antibody conjugated to APC. Cells were analyzed via flow cytometry (Fig 5A). S7-Aerolysin showed a somewhat small but statistically significant increase in the APC/GFP ratio compared to the induced control, 6.04 vs 4.80 respectively, while S7 and the WDV-Aerolysin control did not show a statistically significant difference (Fig 5B). Additionally, we also assessed the shifts in the cell surface anti-Flag-APC vs. GFP cell populations. We initially hypothesized that we would either see a decrease in APC signal or a decrease in both APC and GFP of the S7-Aerolysin population. However, we observed a consistent level of cell surface anti-FLAG-APC when compared to the induced control but a decrease in the GFP positive population (Fig 5C). When quantified, the percentage of the total cell population that falls within this lower GFP area was nearly 20% for the S7-Aerolysin treatment, but around 2-3% for all other treatment conditions (Fig 5D). These data demonstrate that neither S7 nor S7-Aerolysin affect the absolute amount of FLAG-N1-GFP reporter protein localized to the cell surface, thus the inhibition of Notch signaling is not due prevention of trafficking or removal of Notch protein from the plasma membrane.

**Figure 5.**
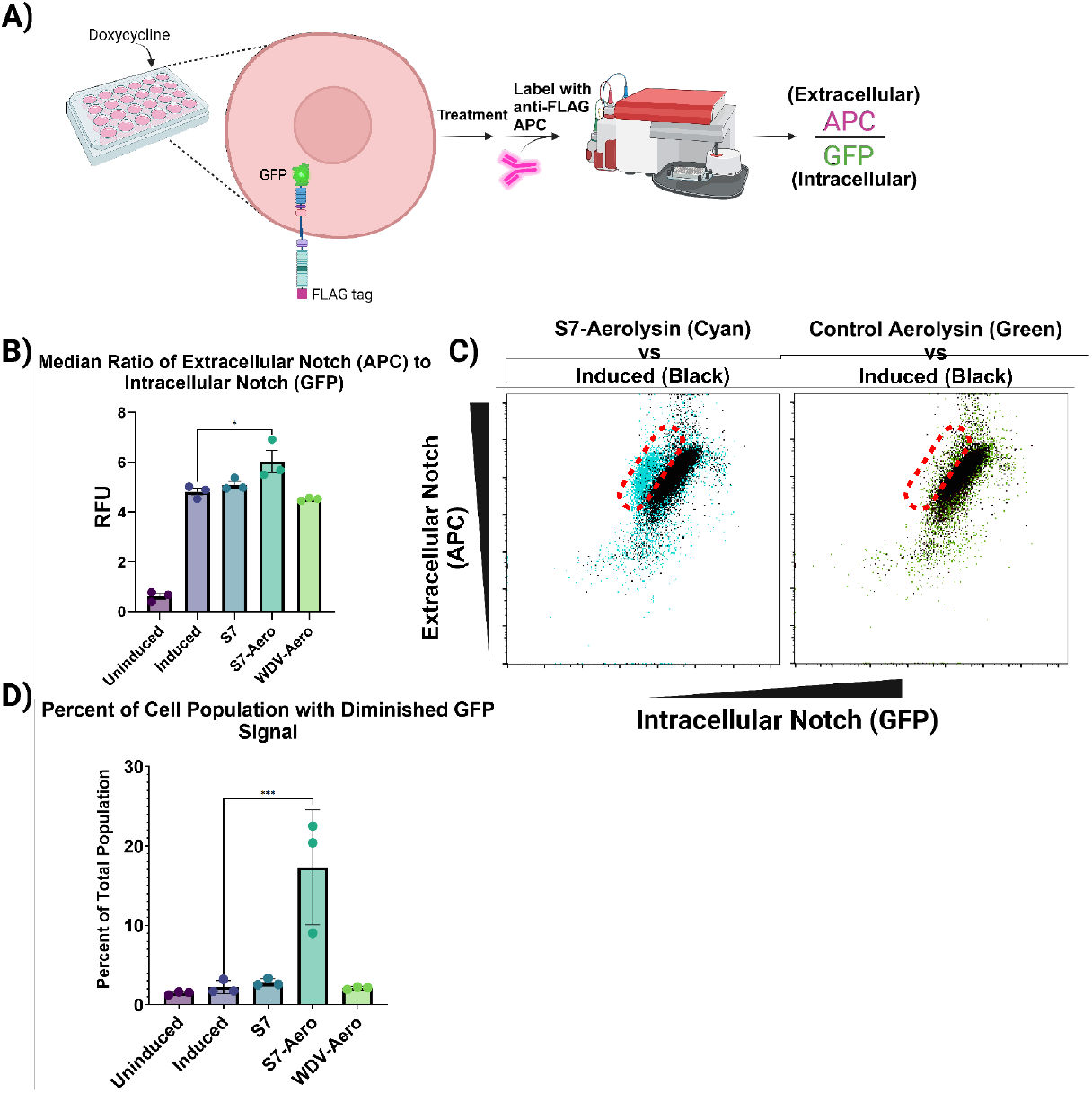
S7-Aerolsyin alters cell surface staining populations of Notch1. (A) Cartoon schematic of the experiment showing the doxycycline induction of the Notch1 receptor with a FLAG tag on the extracellular side, and a GFP fused to the C-terminus of the intracellular side. Cells are treated and then stained with an Anti-FLAG-APC antibody to stain Notch at the cell surface. Cells are then analyzed on flow, and the APC signal is normalized to the overall GFP signal to control for the total amount of Notch in the cell vs Notch just on the cell-surface. (B) Quantification of the Median RFU ratio between APC signal and GFP signal(n=3). Example dot plot of cells gated for APC and GFP signal, with the dotted red circle highlighting the shifted cell population (represented as colored dots) vs the induced population (represented as black dots). (D) Quantitation of the percent of total cell population that resides within the dotted red circle from panel C.

### S7-Aerolysin inhibits cell-cycle progression of HPB-ALL

Given the data demonstrating that S7 acts to inhibit Notch signaling, we wanted to determine if treating cancer cells with S7 or S7-Aerolysin could inhibit cell proliferation, as other inhibitors have been shown to do. To assess this we chose to compare two T-cell acute lymphoblastic leukemia (T-ALL) lines, Jurkat cells and HPB-ALL cells. HPB-ALL is a T-Cell cancer line which has mutations in the Notch1 heterodimerization domain and PEST domain leading to activation of Notch signaling(^27^ and is sensitive to treatment with Gamma-secretase inhibitors (GSI), which inhibit Notch signaling. Jurkat cells are a T-ALL cell line which do have aberrant Notch signaling, but are not sensitive to GSI treatment^28^. It has also been shown that T-ALL cancer lines which are sensitive to inhibition of Notch signaling via GSI will accumulate cells in the G0/G1 phase due to cell cycle blockade^28,29^. Due to this, we compared Jurkat cells against HPB-ALL cells with treatments of S7 and S7-Aerolysin cultured for 2 - 3 days and then fixed. The percentage of cells in G0/G1 phase was measured using flow-cytometry. As expected. treatment with compound E, a gamma-secretase inhibitor, significantly increased the number of HPB-ALL cells in G0/G1 phase but did not have a statistically significant effect on Jurkat cells (Fig 6 A and B), consistent with Notch insensitivity. Interestingly, treatment with S7 did not have a statistically significant effect on HPB-ALL cell-cycle. However, treatment with S7-Aerolysin did show a statistically significant increase in cells in G0/G1, with the non-S7 Aerolysin control showing no effect (Fig 6 A and B). Additionally, neither S7 or S7-Aerolysin showed an effect on the percentage of G0/G1 cells (Fig 6 A and B). Taken together these data show that while S7 treatment was not sufficient to inhibit cell-cycle progression of HPB-ALL cells, S7-Aerolsyin was able to effectively increase the percentage of G0/G1 cells. This likely results from the increased potency and localization of S7 to the cell membrane with the Aerolysin construct.

**Figure 6.**
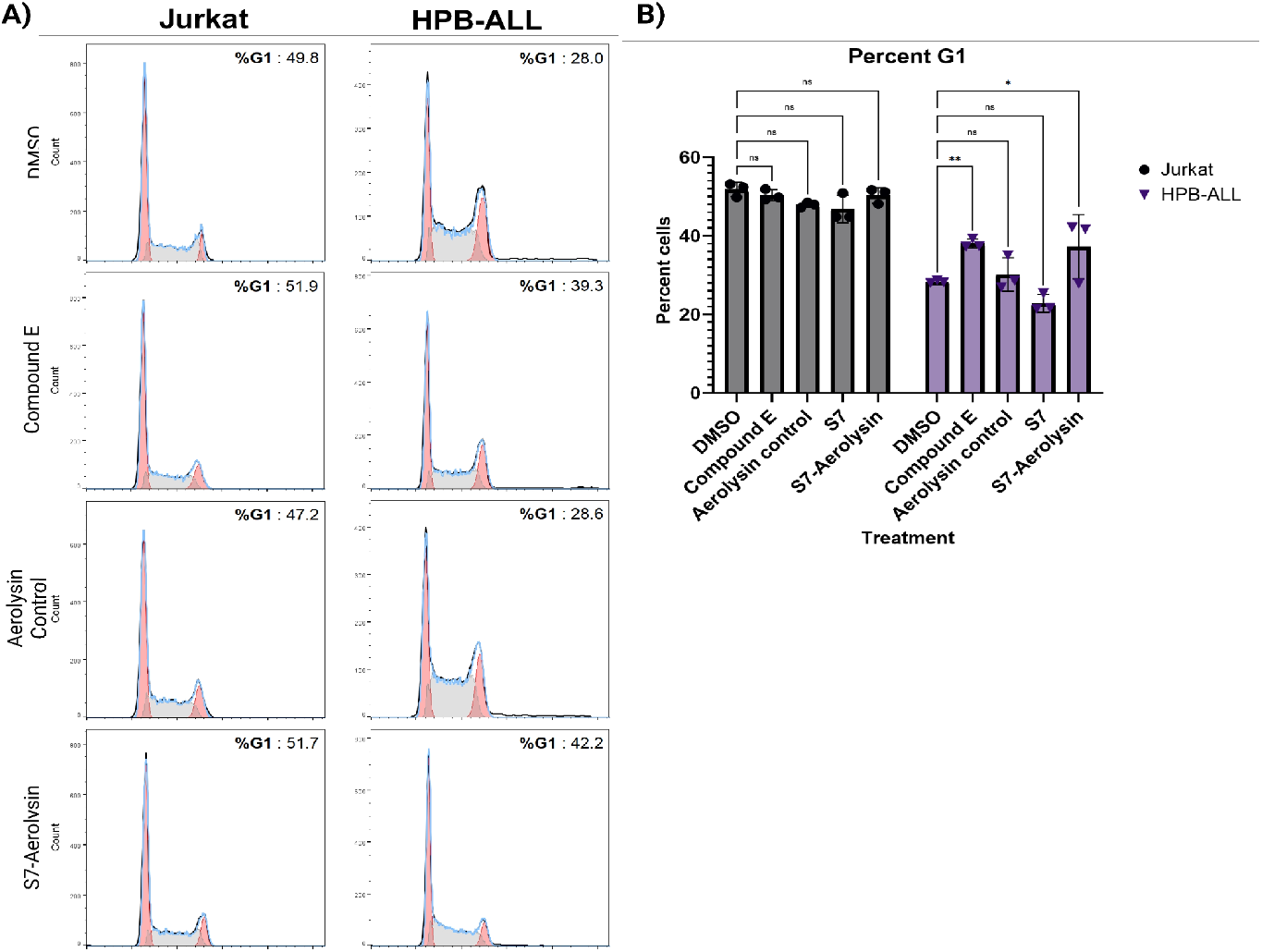
S7-Aerolysin inhibits the proliferation of T-ALL cell line sensitive to Notch-inhibition but does not alter proliferation of Notch-insensitive cell line. (A) DNA staining and cell cycle analysis with Jurkate cells (Notch insensitive) vs HPB-ALL cells (Notch sensitive) with the indicated treatments. (B) Quantification of the percentage of cells in the G1-phase with the various treatments.

## Discussion and Conclusions

In this study, we developed a nanobody inhibitor that binds to the NRR of Notch. Despite modest binding affinity, the nanobody inhibits downstream Notch signaling and modulates terminal differentiation of hematopoietic stem cell progenitors, a precursor stage to T-cell development. This is an interesting finding in itself, as current inhibitory antibodies bind epitopes on both the Lin12/Notch Repeat (LNR) and heterodimerization domain (HD) that are proposed to clamp the two domains in the autoinhibited conformation to prevent signaling^12^. Given the small size of nanobodies, it is unlikely that nanobodies would be able to span the distance between these two domains and inhibit in the same manner. Indeed, results from our cross-linking mass spectrometry experiments suggest a binding site distinct from current epitope-mapped inhibitory antibodies (SuppTable 1, SuppFig 1C), revealing that the NRR structure is susceptible to other modes of allosteric control.

There is also great interest in developing modulators that act as agonists of Notch signaling. These strategies primarily rely on using a protein scaffold upon which Notch ligands or binders are attached^30^. However, an antibody-based binder to the Notch receptor has yet to be developed that is also capable of activating the Notch receptor. This study also provides proof of concept for a workflow to identify allosteric modulators that might serve to activate the receptor.

The cell surface proteolysis that Notch undergoes during its activation is not a mechanism unique to the Notch receptor. In fact, there are hundreds of diverse cell surface proteins that use proteolysis as a mechanism of regulation^31,32^. This study demonstrates that nanobodies are capable of modulating cell-surface proteolysis, and it is likely that nanobodies could be applied to other cell-surface proteins which undergo proteolysis in order to modulate them. Their smaller size and the nature of their antigen binding loops allows them to reach epitopes that might be inaccessible to traditional antibodies^33,34^. Nanobodies that modulate the conformational states of other receptors, including RTKs such as the HER family, immune checkpoint inhibitors (such as PD-1/PD-L1), and protease-associated signaling pathways (MMPs, ADAMs), could lead to novel therapeutic interventions.

We also demonstrated that fusing the nanobody to a toxin derived membrane-associating domain, called Aerolysin, greatly enhanced its inhibitory properties with over a 60-fold decrease in the IC50 of the S7-Aerolysin fusion compared to the S7 nanobody alone (Fig. 5B). We initially hypothesized that this membrane association would simply localize the nanobody in close proximity to its binding epitope and effectively increase its cell surface concentration. However, our flow cytometry analysis of Notch intracellular and extracellular domains alluded to the nanobody-Aerolysin fusion uniquely affecting degradation/trafficking of cell surface Notch. Currently the exact mechanism behind this is not understood, but will be the subject of further studies. More broadly, this strategy of localizing antibodies to cell-surface proteins closer to their surface epitope may allow antibody modulators to generally be used at lower doses. Generally the work presented here provides evidence that Aerolysin could be used as a platform in conjunction with antibodies that may have suboptimal affinities, but unique or desired effects, to still be practical options as research tools.

Lastly, one other aspect of this work that is worth noting is the demonstration that a moderate to low affinity binder requiring a high concentration to have a measurable effect can have its low potency overcome with a fusion to a higher affinity binding partner. One of the major challenges in developing therapeutics that rely on antigen binders with a modulatory effect is also engineering in cell or tissue type specificity. Though the target antigen will often have some level of dysregulation in the disease state that can be exploited, the target of interest is also often still present in healthy cells and tissues leading to off-target effects. One could imagine that this strategy, having a low-affinity modulator paired with a high affinity binder that is cell/tissue-type specific, could be a general methodology to reduce off-target effects while maintaining a therapeutically relevant level of potency, which would be of significant impact within both the realm of research as well as therapeutics.

## Acknowledgements

We would like to acknowledge funding from R33CA272331 (WRG, JMM, JA), R35GM119483 (WRG), and T32GM008347 (ALL). We would like to thank Li Zhang for his help constructing the nanobody library.

