## Supplementary information for "Single-chain nanobody inhibition of Notch and avidity enhancement utilizing the β-pore forming toxin Aerolysin"

| **Table 1. Identified nanobody-NRR crosslinked peptides from CL-MS** | | | |
| --- | --- | --- | --- |
| Nanobody | Crosslinked nanobody peptide | Crosslinked NRR peptide | XlinkX score |
| α-GFP | QAPG[**K**]ER | VLHTNVVF[**K**]R | 65.67 |
| α-GFP | SSYEDSV[**K**]GR | VLHTNVVF[**K**]R | 197.71 |
| S7 | TSYADSV[**K**]GR | AAEGWAAPDALLGQV[**K**]ASLLPGGSEGGRR | 81.02 |
| S7 | [**A**]QVQLVESGGGLVQAGGSLR | VLHTNVVF[**K**]R | 140.63 |
| S7 | QAPG[**K**]ER | VLHTNVVF[**K**]R | 13.58 |
| S7 | TSYADSV[**K**]GR | VLHTNVVF[**K**]R | 241.61 |

Sup. Table 1: **Identified** **nanobody-NRR crosslinks.** Peptides identified from Crosslink-Masspec with associated XlinkX confidence scores.

Sup. Fig1: **Crosslink-MS suggests S7 binds near the Furin loop of the Notch1 NRR.** (A) Cartoon schematic of Cross-linking workflow. NRR and nanobody are co-incubated in the presence of DSSO. Protein is purified out of solution and then analyzed by Mass spectrometry for DSSO-crosslinked peptides. (B) SDS-PAGE gels of purified protein used for cross-linking experiments as well as samples from cross-linked solutions. (C) Notch1 NRR with S7 Cross-linked amino acids labeled in yellow. LNRs A, B and C are colored light pink, pink and red respectively. The HD domain is colored in cyan and the furin loop is colored in black.

Sup. Fig 2:Spectra associated with NRR-ɑ-GFP crosslinked peptide from nanobody-NRR CL-MS. (A) proteins associated with either α (NRR) or β (⍺-GFP-Nanobody) labels by software as well as the identified peptide sequence with the crosslinked residue denoted by |. (B) MS2-spectra identifying the peaks corresponding to crosslinked peptides, the α peptide is labeled red, the β peptide is labeled in blue. (C) MS2-spectra prior to CID. (D) MS3-spectra for the α peptide with fragment ions identified in red or orange.

Sup. Fig 3: **Spectra associated with NRR-S7 crosslinked peptide from nanobody-NRR CL-MS**. (A) proteins associated with either α (S7) or β (NRR) labels by software as well as the identified peptide sequence with the crosslinked residue denoted by |. (B) MS2-spectra identifying the peaks corresponding to crosslinked peptides, the α peptide is labeled red, the β peptide is labeled in blue. (C) MS2-spectra prior to CID. (D) MS3-spectra for the α or β peptide with fragment ions identified in red and orange or blue and cyan respectively.

Sup. Fig 4:**Spectra associated with NRR-S7 crosslinked peptide from nanobody-NRR CL-MS**. (A) proteins associated with either α (S7) or β (NRR) labels by software as well as the identified peptide sequence with the crosslinked residue denoted by |. (B) MS2-spectra identifying the peaks corresponding to crosslinked peptides, the α peptide is labeled red, the β peptide is labeled in blue. (C) MS2-spectra prior to CID. (D) MS3-spectra for the α or β peptide with fragment ions identified in red and orange or blue and cyan respectively.

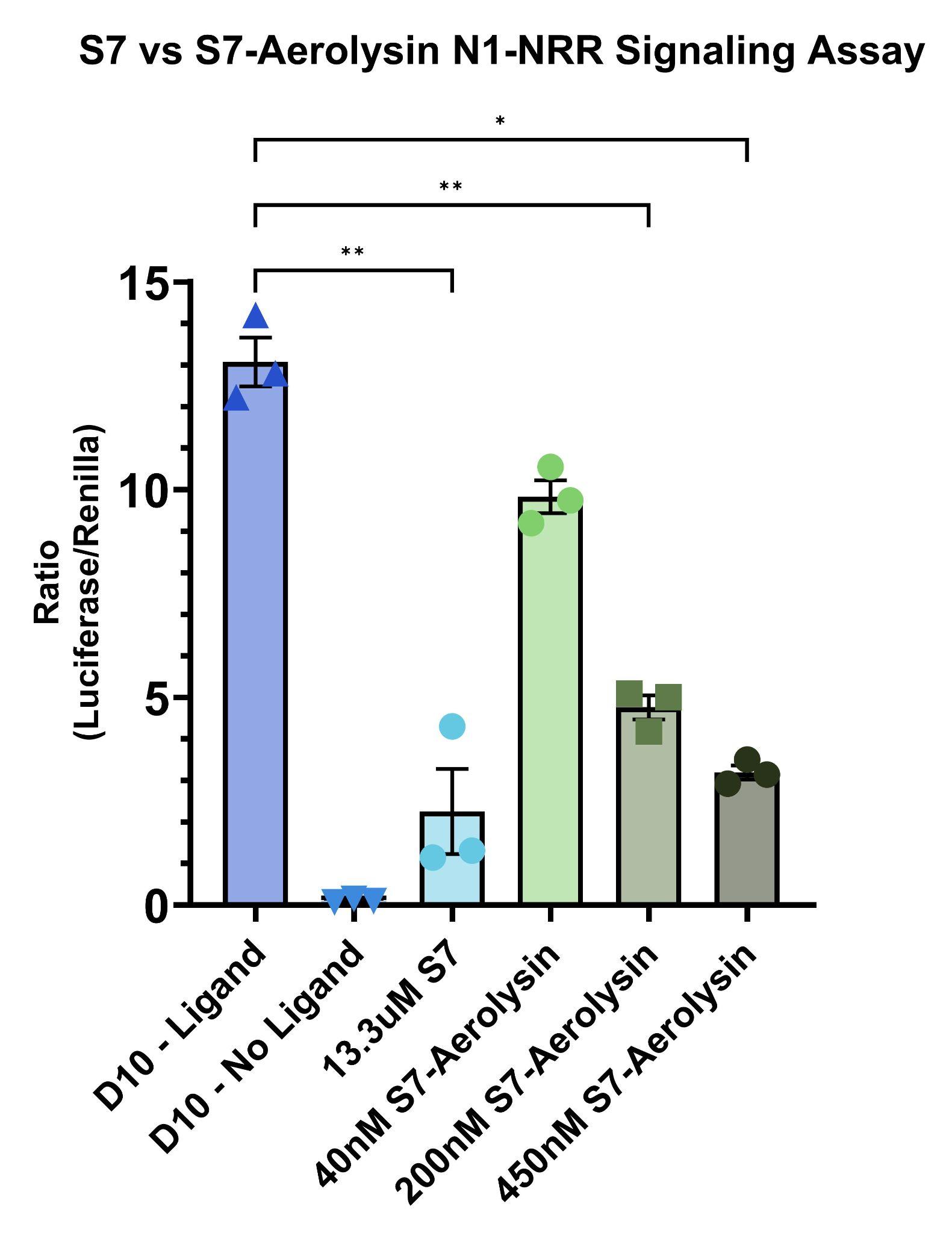

Sup. Fig 5: **S7 vs select concentrations of S7-Aerolysin in signaling assay demonstrating increased efficacy of S7-Aerolysin.** (A) The response of the Notch1 reporter of activation in a cell-based signaling assay to treatment with either S7 or S7-Aerolysin at different concentrations. 200nM and 400nM treatments of S7-Aerolysin demonstrated statistically significant inhibition compared to the positive ligand control while 40nM S7-Aerolysin qualitatively reduced signal but was not statistically significant.

Sup. Fig 6: **S7 and S7-Aerolysin treatments attenuate Notch1-reporter line response to DLL4 ligand in a manner consistent with an inhibitor of Notch signaling.** (A) Cartoon schematic of the experimental setup. Coats are plated with a titration of DLL4 ligand and cells reverse transfected with the Notch1-reporter system are then plated and treated with either 10uM S7 or 250nM S7-Aerolysin. (B) Cells treated with 10uM S7 show a rightward shift in the dose response, indicating less sensitivity to DLL4. Cells treated with 250nM S7-Aerolysin show a similar decrease in response to DLL4 ligand.
